# Isolation and Purification of Active Antimicrobial Peptides from *Hermetia illucens* L., and Its Effects on CNE2 Cells

**DOI:** 10.1101/353367

**Authors:** Zhong Tian, Qun Feng, Hongxia Sun, Ye Liao, Lianfeng Du, Rui Yang, Xiaofei Li, Yufeng Yang, Qiang Xia

## Abstract

Active antimicrobial peptide HI-3 was isolated and purified from the 5^th^ instar larvae of *Hermetia illucens* L., and its effects on proliferation, apoptosis and migration of nasopharyngeal carcinoma (CNE2) cells were investigated. The expressions of telomerase reverse transcriptase (hTERT) in CNE2 cells were also studied *in vitro* to elucidate the mechanism involved in the action of HI-3 on CNE2 cells. Results showed that three fractions (HI-1, HI-2, HI-3) were isolated from the hemolymph of *H. illucens* larvae. After purified by RP-HPLC, only HI-3 showed the inhibitory activities to four strains of bacteria. It was also showed that HI-3 could effectively inhibit the proliferation of CNE2 cells in a dose- and time-dependent manner. Apoptosis of CNE2 cells was observed in the treatment with 160 μg/ml HI-3, and the early apoptosis rate up to 27.59 ± 1.14%. However, no significantly inhibitory effects and apoptosis were found on human umbilical vein endothelial cells (HUV-C). Moreover, HI-3 could significantly reduce the migration ability of CNE2 cells when compared with that of the control. On the other hand, the levels of mRNA and protein of hTERT in the HI-3 treatment were all significantly lower than that of the control. Results indicated that HI-3 could inhibit the proliferation of CNE2 cells and induce the apoptosis of CNE2 cells by down-regulating the telomerase activity in CNE2 cells, while no obvious effect was occurred on HUV-C. It inferred that HI-3 is a potential anti-tumor drug with low toxicity to normal cells.

**Summary Statement:**

1. An active antimicrobial peptide HI-3 was isolated and purified.
2. Inhibitory proliferation of CNE2 cells, but no effect on normal cells.
3. A potential antitumoral drug.

## 1. Introduction

More and more attentions were paid on the nasopharyngeal carcinoma (NPC) for its high incidence rates in Asia. Data from IARC showed that the incidence rates of NPC were up to 71.02% for men, which was nearly 2.45 times higher than that of women in Asian countries, especially in the country of Malaysia, Singapore, Indonesia and China (Mahdavifar et al., 2016). Currently, the traditional surgery combined with chemoradiotherapy was widely applied during its therapy. However, the radiotherapy and chemotherapy would inevitably damage the normal cells and the patient’s immune systems (Ng and Lee, 2017), which more or less lead to the toxicity and severe effects in some tissues and organs (Thangavel et al., 2016). And the strong invasion ability of NPC cells and its inclination to distal metastasis made its treatments become more difficult (Jiang et al., 2014; Yang et al., 2013). Therefore, looking for a new, safe and effective anticancer agent for treatment of NPC has become a hot research direction worldwide.

Antimicrobial peptides (AMPs), the small molecule peptides that were firstly discovered by Boman research team, were found to have anti-bacterial, anti-viral, and anti-fungal properties. It was reported that AMPs could act rapidly to combat the invasion of potential pathogens, and thus serve to limit the extent of infection prior to activation of the adaptive immune response (Wanmakok et al., 2018). Furthermore, some of these peptides were found to display a wide range of biological activities and have been shown to exert antitumoral activity (Cerón et al., 2010). Nowadays, AMPs are considered as a potential alternative for the treatment of emerging drug-resistant infections and cancer (Riedl et al., 2011). The detailed mechanisms involved in antitumoral activities of AMPs were also gradually established. Studies showed that AMPs might disrupt tumor cell membranes integrity and act on mitochondria or DNA to exert its antitumoral activity (Lemeshko, 2013). It was also reported that AMPs perform its anticancer effects by regulating the organisms’ immune system. Studies have indicated that the organisms’ immune defenses were enhanced by inducing dendritic cell aggregation and increasing the expression of lymphocytes after AMPs treatment, and the development of cancer was also inhibited (Huang et al., 2015).

In recent years, it was revealed that the telomerase might also be involved in the antitumoral mechanisms of AMPs. In normal cells, the expressions of telomerase were extremely low, or even none. However, their expression levels were significantly increased in malignant proliferative cells to protect their chromosomes and to promote the cells to proliferate indefinitely (Li et al., 2014; Ale-Agha et al., 2014). Generally, the main structure of telomerase includes telomerase reverse transcriptase (TERT) and telomerase-associated proteins (Blackburn et al., 2006). TERT, as the catalytic subunit of telomerase, plays a key role in telomerase synthesis and telomerase activity maintenance, especially in promoting the cells carcinogenesis (Borah et al., 2015). It was reported that higher levels of TERT mRNA were expressed in lung cancer-derived human cell line H1299, while lower levels in those of normal cells (Bodnar and Wright, 1998). Study also found that plasma TERT mRNA levels were significantly higher in the patients with liver cancer and prostate cancer than those in the normal group (El-Mazny et al., 2014), and the levels of TERT mRNA were closely related to various clinical and pathological conditions (Kang et al., 2013). Similar results were also reported by FU et al. (2015), which found that plasma hTERT mRNA levels in peripheral blood were significantly higher in patients with NPC than those in the normal groups through the quantitative detection of telomerase hTERT mRNA. In addition, the levels of telomerase hTERT mRNA were also been shown to be up-regulated, followed by the G1/S blockade and chemotherapeutic drug resistance under hypoxia in NPC. Until the hypoxia reversed, the telomerase activities were decreased, and chemo-resistance was improved (Shi et al., 2015). From these studies, it was inferred that the telomerase activity was closely related to the occurrence and development of NPC.

The black solider fly, *Hermetia illucens* L. is considered as a beneficial resource insect in Diptera family. Its larvae are reported as feeding on immense variety of organic material, and have been used in small-scale waste management purpose to reduce manure accumulations in confined animal feeding operations and the accumulations of other wastes. The habit of *H. illucens* larvae, which is extremely rich in various microorganisms including many pathogenic ones, implies that the immune system of this insect functions works very efficiently (Park et al., 2014). Since its first report in 2013 by our research group (Xia et al., 2013), AMPs of this insect, which are essential components of innate immune system of *H. illucens* larvae, have been paid more and more attentions and been considered to have potential applications in the medicine fields (Choi and Jiang, 2014; Park et al., 2015; Vogel et al., 2018). AMPs of *H. illucens* have showed good antibacterial activity for gram-positive bacteria, gram-negative bacteria and fungi (Elhag et al., 2017). However, no research was focused on their anti-cancer activity and related mechanisms. Therefore, in the present study, the AMPs were isolated and purified from the hemolymph of *H. illucens* larvae, and the active components HI-3 were screened out to investigate its effects on the proliferation, apoptosis and migration of nasopharyngeal carcinoma (CNE2) cells. Furthermore, the expressions of telomerase in CNE2 cells were also studied to explore the possible mechanism of HI-3 action. We hope the results would provide theoretical basis for the research and development of the *H. illucens* AMPs as a potential antitumoral agent.

## 2 Materials and methods

### 2.1 Insects, bacteria and cells

*H. illucens* L. was provided by Zhuhai Modern Agriculture Development Center. The larvae were feed with artificial feed (wheat bran: corn flour: chick feed=5: 3: 2) and kept in a greenhouse of 27, RH 80%, photoperiod 14 L: 10 D.

The four strains of bacteria *Staphylococcus aureus, Escherichia coli, Bacillus subtilis*, and *Enterobacter aerogenes* were obtained from Immunology and Pathogenic Biology Laboratory of Zunyi Medical University.

Standard strains of human CNE2 cells and human umbilical vein endothelial cells (HUV-C) were purchased from Shanghai ATCC Cell Bank of Chinese Academy of Sciences.

Tricine-SDS PAGE gel kit was purchased from Sigma; CCK-8 kit, Hoechst33342 staining kit, and Annexin V-FITC/PI double staining kit were purchased from Japan Tongren Institute of Chemistry; RT-PCR telomerase detection kit was purchased from Nanjing Kaiji Biological Co., Ltd.; hTERT antibody was purchased from ABclonal Company.

Vertical electrophoresis (Mini-PROTEAN Tetra), Horizontal electrophoresis(YCP-31DN), Automatic microplate reader(Elx-800), RT-PCR (621BR07707) were from Bio-Rad; RP-HPLC (LC-C8002) was from Shimadzu Corporation; Inverted fluorescence microscopy (IX71) was from OLYMPUS Corporation; Flow cytometry (FACS Calibur) was from Beckman Coulter Corporation.

### 2.2 Induction and Extraction AMPs from the larvae of *H. illucens*

According to the method mentioned in Xia et al (2013), the 20,000 fifth instar larvae of *H. illucens* were selected for induction. The larvae in the induction group were received abdomen acupuncture treated with *Staphylococcus aureus* (A260=2.4), while those in the control group were acupunctured with culture medium. After 24 h treatment, the hemolymph of larvae in the induction and control group was collected into Eppendorf centrifuge tubes and centrifuged at 12,000 rpm for 10 min at 4°C. Then, the supernatant was transferred into a 10 kD ultrafiltration centrifuge tube and centrifuged at 12,000 rpm for 15 min at 4 °C. The ultrafiltrate was again transferred to a 3 kD ultrafiltration centrifuge tube and centrifuged under the same conditions. The retention section, which is AMP crude extract, was freeze-dried in vacuo and stored in a refrigerator at −80 for use.

### 2.3 Purification of AMPs from crude extract

The crude extract of AMPs was separated by RP-HPLC system. The column was the type of LC-C8002, 4.6 mm × 250 mm, 10 μm, and column temperature at 40. The mobile phase was 0.1% (v/v) TFA (phase A) and acetonitrile with 0.1% TFA (phase B) at a flow rate of 0.5 ml/min. UV detector wavelength was set at 214 nm. According to the retention time of the peak, the components corresponding to each peak were collected.

Purity of HI-3 was also detected by RP-HPLC. And the chromatographic conditions were same as mentioned above 0.5 ml/min, detection wavelength 214 nm, temperature 40 .

### 2.4 Antibacterial activity assays

Four strains of bacteria, *Staphylococcus aureus, Escherichia coli, Bacillus subtilis* and *Enterobacter aerogenes*, were formulated into bacterial liquid at the concentration of 3.2 ×10^12^/ml, separately. Then, 50 μl of each bacterial liquid was evenly coated in the LB solid medium. When the bacterial liquid was dry, the sterile filter paper with a diameter of 6mm, which were treated with 25 μl of Gentamicin sulfate (as positive control), saline (as negative control) and purified AMPs (named as HI-1, HI-2, HI-3) separately, were placed on the medium. Four replicates were set. After incubated at 37 °C for 24 h, the size of the inhibition zone was observed and measured (Xia et al., 2013).

The value of MIC (minimum inhibitory concentration) about the component HI-3 with strong antibacterial activity was detected by MH broth dilution method. Gentamicin sulfate was used as positive control. Absorbance value was read at 595 nm to calculate the inhibition rate Y.
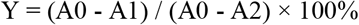

Among them, A0 is the absorbance value in the negative control, A1 is the value in the HI-3 treatment, and A2 is the value in positive control.

MIC was recorded as the minimum inhibitory concentration at which the inhibition rate was not less than 80%.

### 2.5 Estimation of cytotoxicity in CNE2 cells and HUV-C

To test the cell viability after treatment with HI-1, HI-2 and HI-3, CCK-8 kit was run. Cells were plated at a density of 1×10^5^ cells/100 μl/well in 96-well plates for 24 h. Four replicates were set for each treatment. Then, the medium was discarded and replaced by different concentrations (5 μl/ml, 10 μg/ml, 20 μg/ml and 40 μg/ml) of HI-1, HI-2, HI-3, respectively, or medium alone for 24 h and 48 h. This was followed by the addition of 20 l of CCK-8 solution for 2 h at 37 °C in 5% CO_2_ incubator. The optical density was measured spectrophotometrically at 490 nm on a microtiter plate reader. Results were expressed as a percentage of the inhibition rate for viable cells, and values of the medium only group were regarded as negative control.
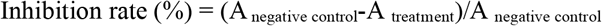

### 2.6 Effects of HI-3 on apoptosis and migration of CNE2 cells

#### 2.6.1 Apoptosis of CNE2 cells

The CNE2 cells and HUV-C were maintained in RPMI-1640 and DMEM medium supplemented with 10% FBS, respectively. Cells were maintained in incubator under a fully humidified atmosphere of 95% room air and 5% CO_2_ at 37. Briefly, the cell density was adjusted to 1×10^5^ cells/ml, and 100 μl of cell suspensions were placed onto 96-well plates. Four replicates were set for each treatment. The medium was removed after 24 h, and HI-3 was added to cell cultures at a final concentration of 40 μg/ml, 80 μg/ml and 160 μg/ml. After incubation for 48 h at 37 in a humidified atmosphere with 5% CO_2_, 5 μl of Hoechst33342, a DNA specific fluorescent dye, which stains the condensed chromatin of apoptotic cells more brightly than the chromatin of normal cells, was added to each well and incubated for 15 minutes at room temperature. Morphological changes in nuclear chromations of cells in control and each treatment were observed with fluorescence microscope, and the influence of HI-3 on the apoptosis rate of CNE2 cells and HUV-C was detected by flow cytometry (FCM) (Yan et al., 2015).

#### 2.6.2 Migration of CNE2 Cells

The HUV-C and CNE2 cells suspension was prepared as the method mentioned in 2.7.1. 2 ml of cell suspension was placed into a 6-well plate and incubated at 37 °C in the incubator with 5% CO_2_. After the cells were fully covered the plate, the cells in each well was perpendicularly streaked with a 200 μl of pipette tip, followed by washing with PBS. Then, 2 ml of HI-3 diluted with 2% FBS and 0.2% colchicine was added to cell cultures at a final concentration of 160 μg/ml, while the negative control received with the same volume of solution without HI-3. After 0 h, 12 h, 24 h and 48 h incubation, the scratches were observed and photographed to calculate the mobility.
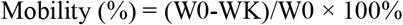

Among them, W0 is the width of scratches at 0 h, WK is the width of scratches at 12 h, 24 h, 48 h, respectively.

### 2.7 RT-PCR analysis of telomerase hTERT gene expression

V RT-PCR was used to analyze gene expression of telomerase hTERT in CNE2 cells treated with 160 μg/ml of HI-3 for 48 h. And the cells without treatment were set as control. The primers used for hTERT were upstream 5′-GCCGATTGTGAACATGGACTACG-3′ and downstream 5′-GCTCGTAGTTGAGCACGCTGAA-3′. Primer pair for GAPDH (upstream 5′-CATCTTCTTTGCGTCGCCA-3′; downstream ′5-TTAAAAGCAGCCCTGGTGACC-3 ′) was used as the reference gene. Reaction system involved each 1 μl of primers of upstream and downstream, 2 μl of template RNA, 2.75 μl of RT-PCR MIX (including reverse transcriptase, Taq DNA polymerase and RNase inhibitor), 25 μl of 2× reaction buffer, and RNase free water which was added to a total volume of 50 μl. The samples were denatured at 94 °C for 5 min, followed by a 40 cycles reaction: 94 °C for 30 s, 60 °C for 30 s, 72 °C for 1 min; and by the extension at 72 °C for 10 min. The 10 μl of PCR products were separated by 1% agarose gel electrophoresis, and the expressions of hTERT gene were analyzed by 2^−△△ Ct^ method using GAPDH as internal reference gene.

### 2.8 Western blotting analysis of the telomerase hTERT protein expression

Western blotting for detecting telomerase hTERT was performed after CNE2 cells were treated with HI-3 at the final concentration of 160 μg/ml for 48 h. Briefly, the cells were harvested and the cellular protein were extracted by homogenization in 100 μl lysate on ice for 30 min. The lysates were then centrifuged at 12,000 rpm for 30 min at 4. The proteins were quantified using the Bradford method, and 30 μg proteins were separated using SDS-PAGE gel and transferred onto PVDF membranes (Millipore, USA). Non-specific binding was prevented using a solution of 5% non-fat dry milk at room temperature for 1 h, and then washed with TBST for 3 times. Blots were incubated with the primary antibody diluted 1: 1000 primary antibody (for hTERT and GAPDH) diluted with 2% BSA in TBS containing 0.1% Tween-20 (TBST) overnight at 4 °C. After washing with TBST for 3 times, the blots were incubated with mouse-derived secondary antibody diluted 1: 3000 with 1% BSA in TBST for 1 h at room temperature. Following additional washing, gray-scale of specific bands was detected with ECL development.

### 2.9 Statistical analysis

Results were expressed as means ± SEM. To analyze the differences between control and treatments, single-factor analysis of variance in SPSS 20.0 software was used. Statistical differences were considered significant at a value of *P* < 0.05.

## 3 Results

### 3.1 Isolation and purification of antimicrobial peptides

The crude extract from the hemolymph of *H. illucens* larvae is clear and transparent, and exhibits white powder after freeze-dried in vacuo. After PR-HPLC analysis, three compounds, HI-1, HI-2 and HI-3, were well separated from the crude extract in the induced group (Figure 1-1)). And the purity of HI-3 was up to 96.1% after purification (Figure 1-2)).

**Figure 1.**
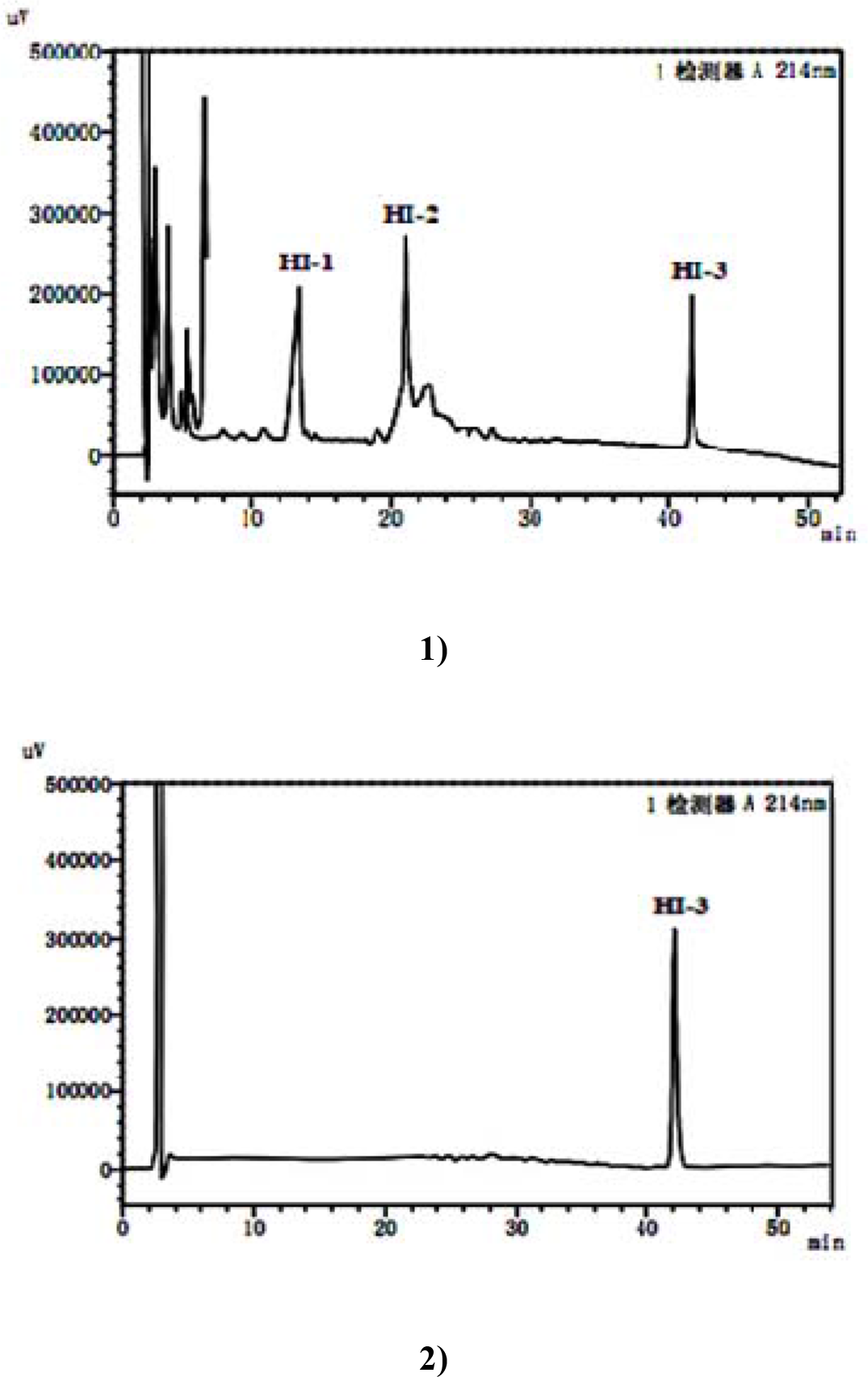
Separation crude extract from the hemolymph of *H. illucens* larvae by RP-HPLC 1) Separation of crude extract; 2) Purity detection of HI-3 by RP-HPLC.

### 3.2 Antibacterial activity of three separated components

As shown in Table 1, only HI-3 showed significant antibacterial activity on the all four strains of bacteria when compared with that in the negative control (*P* < 0.05). Furthermore, the antibacterial activity of HI-3 to *Bacillus subtilis* (10.84 ± 1.00 mm) *Staphylococcus aureus* and *Escherichia coli*, which inhibition zone diameter was 10.84 ± 1.00 mm, was significantly lower than that to *Staphylococcus aureus* (11.79 ± 1.178 mm), *Escherichia coli* (11.83 ± 0.82 mm) and *Enterobacter aerogenes* (11.41 ± 0.78 mm) (*P* < 0.05).

**Table 1.**
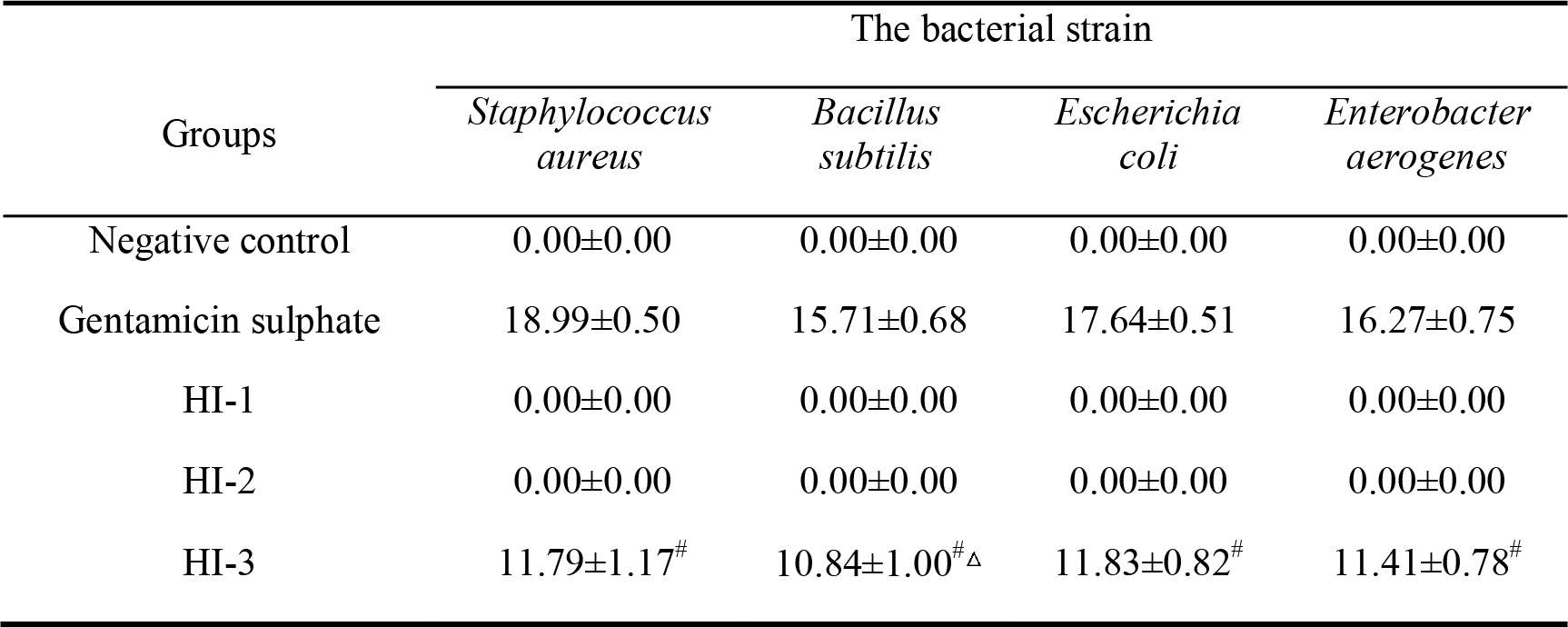
Inhibition zone diameters against the four strains of bacteria after treated with HI-1, HI-2 and HI-3 (n=4, 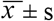, mm) Note: ^#^ *P* < 0.05 indicated there were significant differences in inhibition zone diameters between HI-3 group and negative control. And △ *P* < 0.05 indicated the inhibition zone diameters in the *Bacillus subtilis* treatment were significantly lower when compared with the *Staphylococcus aureus* and *Escherichia coli* treatments in the HI-3 group.

| Groups | The bacterial strain |  |  |  |
| --- | --- | --- | --- | --- |
|  | *Staphylococcus aureus* | *Bacillus subtilis* | *Escherichia coli* | *Enterobacter aerogenes* |
| Negative control | 0.00±0.00 | 0.00±0.00 | 0.00±0.00 | 0.00±0.00 |
| Gentamicin sulphate | 18.99±0.50 | 15.71±0.68 | 17.64±0.51 | 16.27±0.75 |
| HI-1 | 0.00±0.00 | 0.00±0.00 | 0.00±0.00 | 0.00±0.00 |
| HI-2 | 0.00±0.00 | 0.00±0.00 | 0.00±0.00 | 0.00±0.00 |
| HI-3 | 11.79±1.17<sup>#</sup> | 10.84±1.00<sup>#△</sup> | 11.83±0.82<sup>#</sup> | 11.41±0.78<sup>#</sup> |

### 3.3 MIC of HI-3 to the four strains of bacteria

From Figure 2, it was indicated that inhibitory rates of HI-3 against the four types of bacteria were all positively correlated with the concentration of active component HI-3. After analysis, the MICs of HI-3 to *Staphylococcus aureus*, *Bacillus subtilis*, *Escherichia coli* and *Enterobacter aerogenes* were 80 μg/ml, 160 μg/ml, 80 μg/ml and 80 μg/ml, respectively. From these results, it can be seen that the HI-3 was less sensitive to *Bacillus subtilis* than those of the other three bacteria, which was similar to the results of antibacterial activity of HI-3.

**Figure 2.**
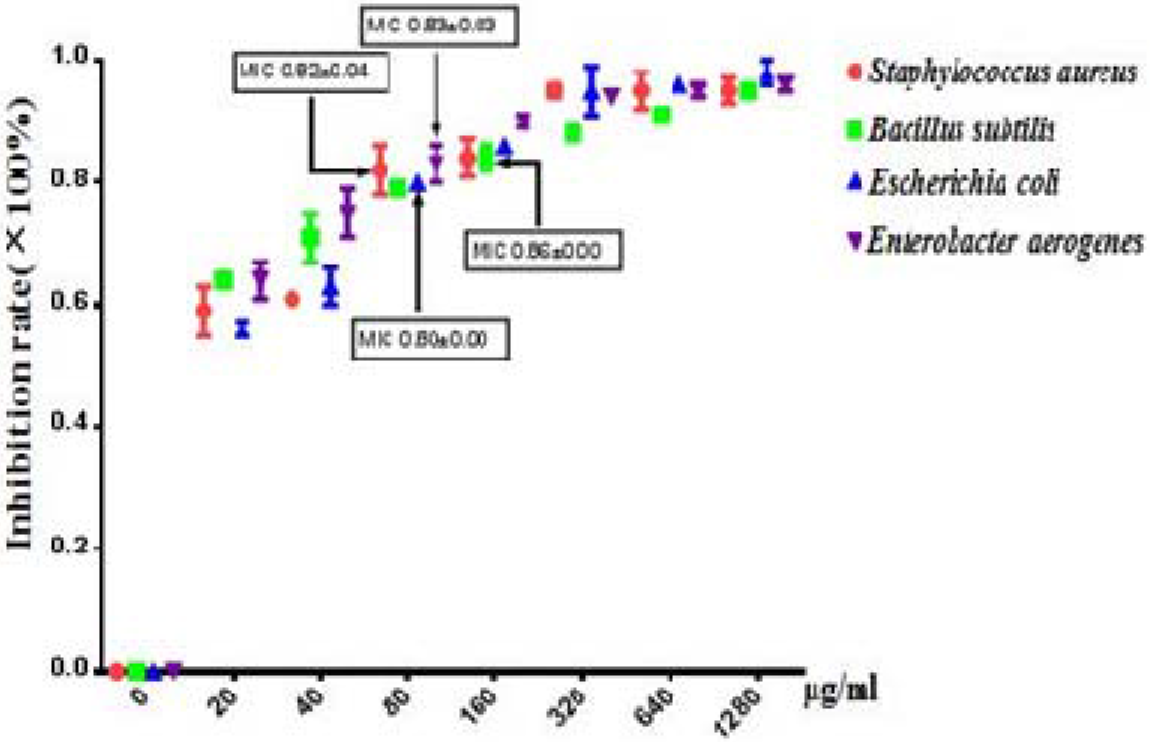
MIC of HI-3 against four strains of bacteria.

### 3.4 HI-3 induces cytotoxicity on CNE2 cells

According to the growth curve of CNE2 cells and HUV-C detected by CCK-8 kit, CNE2 cells entered logarithmic growth phase on the 2nd day, while HUV-C on the 3rd day. Therefore, CNE2 cells after 2 days of culture and HUV-C after 3 days of culture were chosen for follow-up experiments, respectively, to verify the effects of HI-1, HI-2 and HI-3 on the proliferation of the two cells.

The dose- and time-dependent effects of HI-1, HI-2 and HI-3 against CNE2 cells were shown in Table 2. No significant effects on the proliferation of CNE2 cells were found after treatment with HI-1 and HI-2 for 12 h, 24 h and 48 h, respectively. When the concentration of HI-3 was among 40-160 μg/ml, HI-3 exhibited significantly inhibitory activity on the proliferation of CNE2 cells when compared to that in the corresponding HI-1 and HI-2 groups (*P* < 0.05). Furthermore, the inhibitory rate on the proliferation of CNE2 cells also increased with the treatment period of HI-3 at the concentration of 40-160 μg/ml, and the maximal inhibitory rate was recorded in the 160 μg/ml HI-3 after 48 h treatment (Table 2; Figure 3). By contrast, HI-3 treatment had no significant effect on the proliferation of HUV-C (*P*> 0.05) (Data not shown).

**Figure 3.**
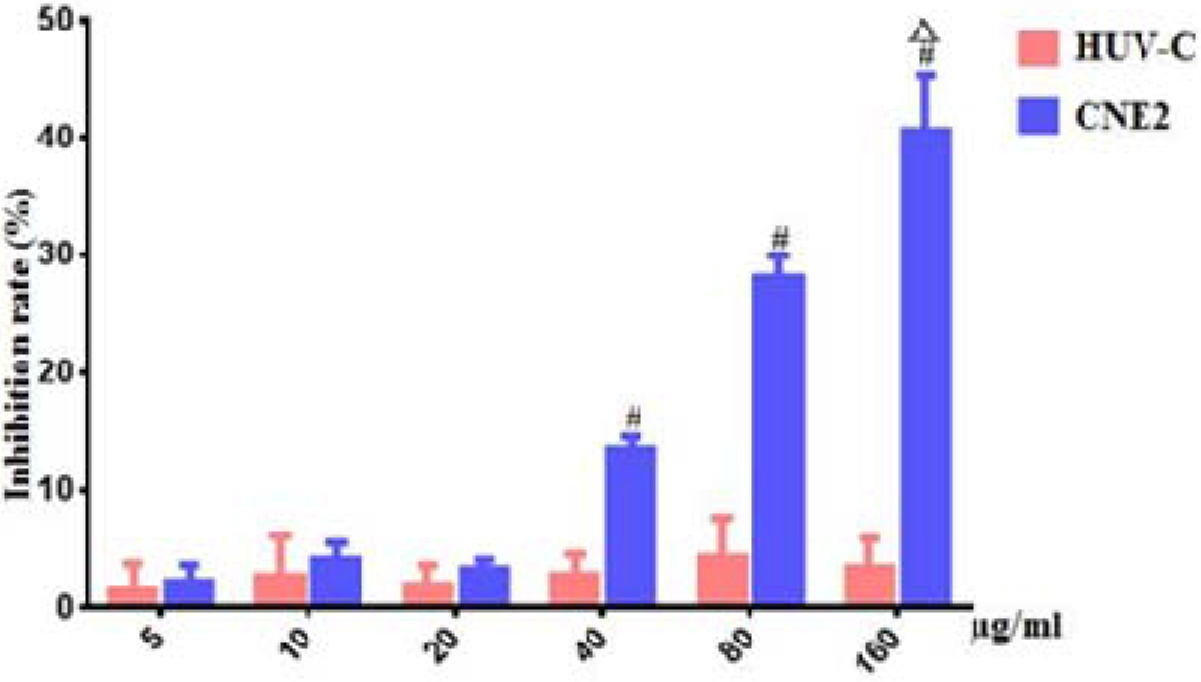
Effects of HI-3 on proliferation inhibition rates of CNE2 Cells and HUV-C after 48 h treatment. Note: # means significant difference between the CNE2 cells group and HUV-C group under the same concentration at the levels of 0.05. △ means this significant difference at the levels of 0.01.

**Table 2.**
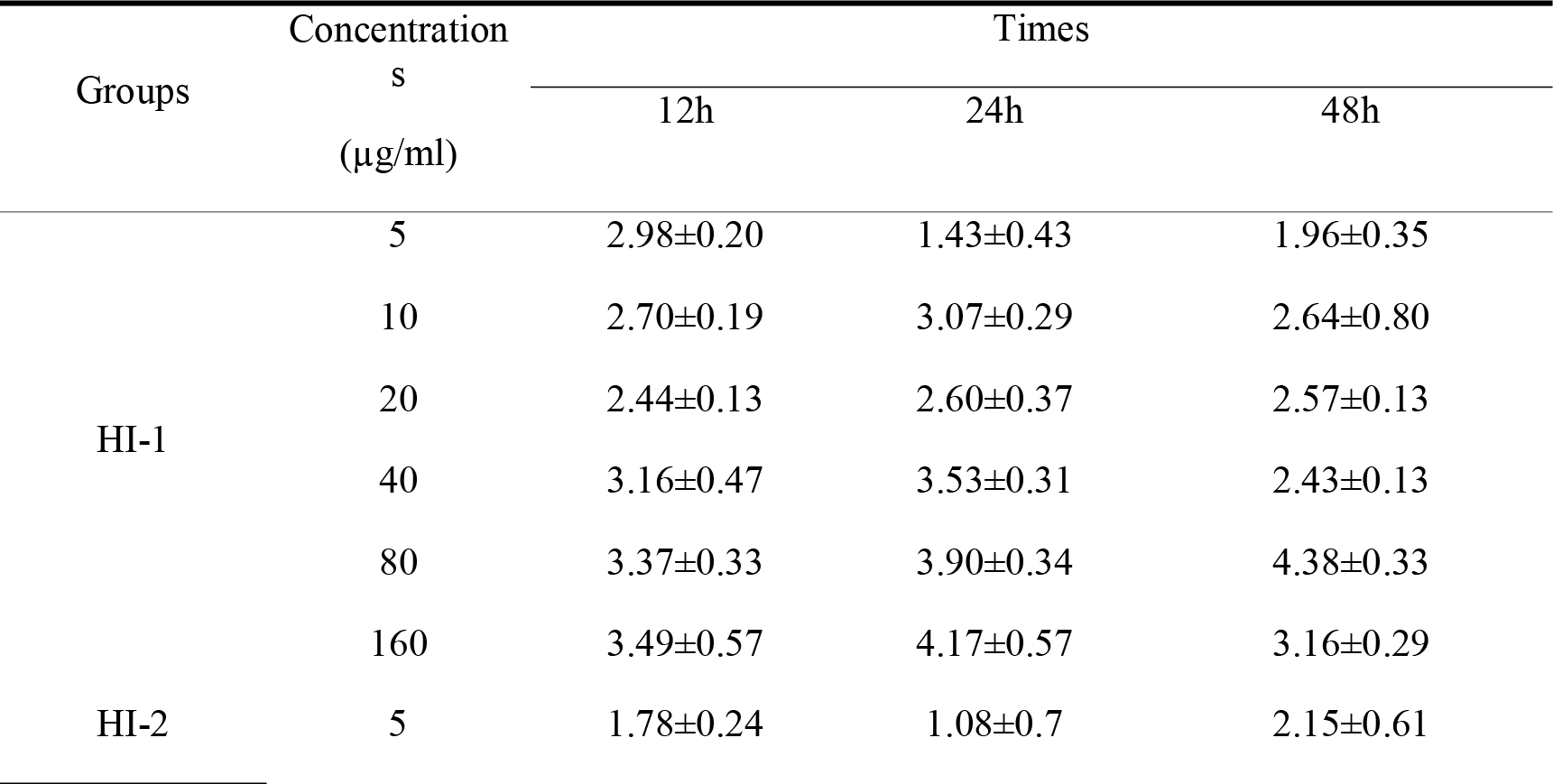
Effects of HI-1, HI-2 and HI-3 on proliferation inhibition rates of CNE2 cell (n=4, 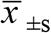, Inhibition rate: %) Note: ^#^ *P* < 0.05 indicated that there were significant differences in the inhibition rates between HI-3 group and HI-1/HI-2 group under the same concentration. △ *P* < 0.05 indicated that there were significant differences in inhibition rates between different time at the same concentration of HI-3 group.

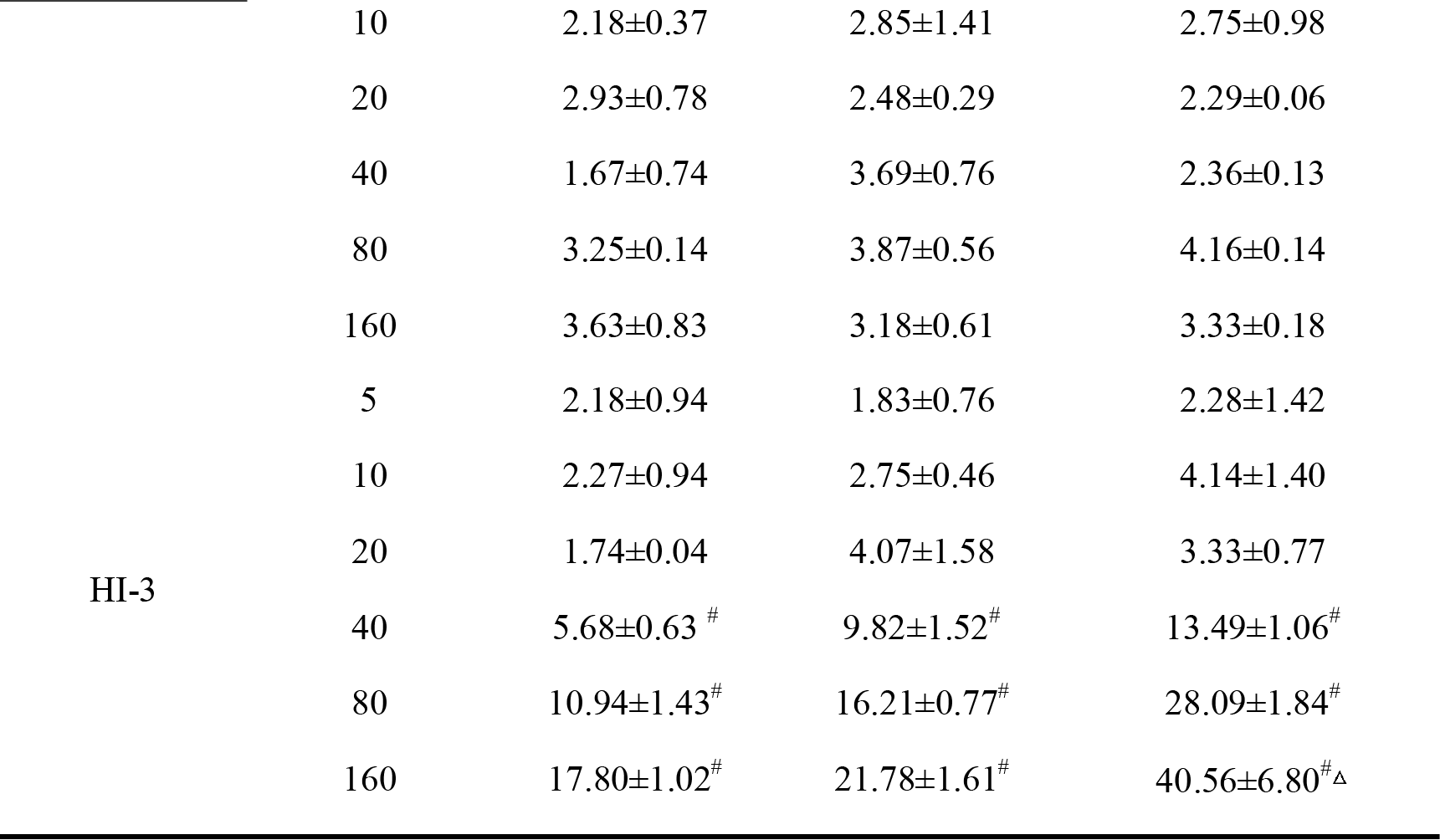

### 3.5 HI-3 induced apoptosis of CNE2 cells

To investigate the mode of action underlying the cytotoxic activity of HI-3 at the concentrations of 40-160 μg/ml, the CNE2 cells and HUV-C were incubated with HI-3 for 48 h for fluorescence microscope and FCM.

Representive micrographs by fluorescence microscope revealed that the CNE2 cells in the control was membrane-intact and light blue, and the emitted fluorescence was weak (Figure 4-1)-A). After 48 h treatment with 40 μg/ml HI-3, increased fluorescence intensity was observed in the CNE2 cells, indicating that HI-3 disrupted the integrity of the plasma membrane, and the symptoms of early apoptosis occurred. However, the fluorescence intensity is still uniform, which meant that the chromosomal DNA was not destroyed. With the increasing HI-3 concentration, the fluorescence intensity in CNE2 cells was further improved, indicating that the permeability of the plasma membrane was gradually increased (Figure 4-1)-B, 4-1)-C). When the concentration of HI-3 was up to 160 μg/ml, the CNE2 cells exhibited the strongest fluorescence intensity, and the number of brightly stained cells with DNA damage was also increased. Furthermore, the typical characteristics of cell apoptosis, such as highly aggregated and fragmented chromatin, and dispersed apoptotic bodies, were appeared. These results exhibited that high concentration of HI-3 induced drastic changes in the cellular morphology (Figure 4-1)-D, arrow).

**Figure 4.**
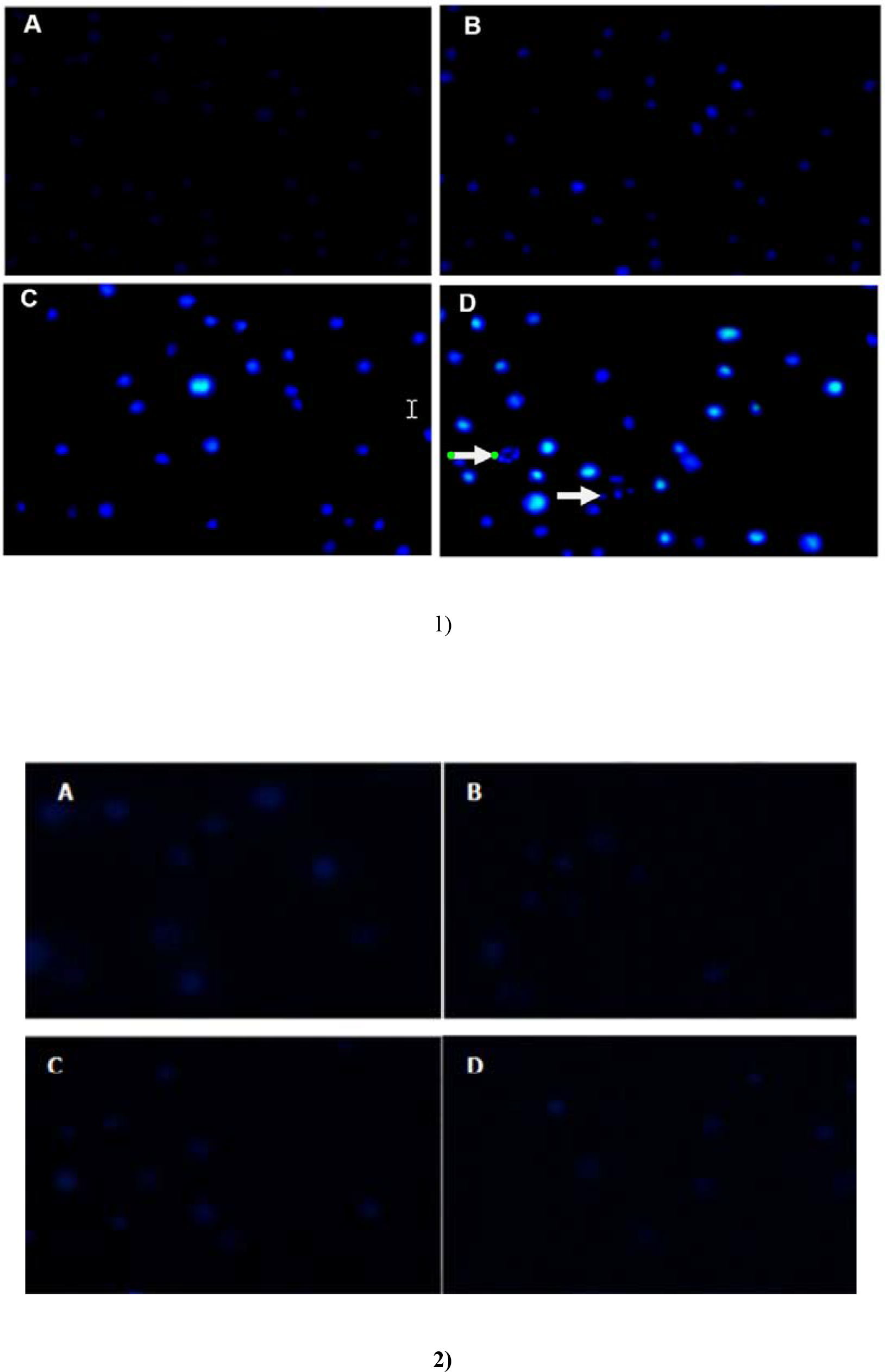
Effects of HI-3 on morphology of CNE2 cells and HUV-C (400 ×) Note:

1. CNE2 cells A: Negative control; B: 40 μg/ml; C: 80μg/ml; D: 160 μg/ml
2. HUV-C A: Negative control; B: 40 μg/ml; C: 80μg/ml; D: 160 μg/ml

By contrast, there were no obvious changes in the morphology of HUV-C after treated with different concentrations of HI-3 for 48 h (Figure 4-2)).

Results from FCM detection showed that exposure to HI-3 at the concentrations of 40-160 μg/ml for 48h caused a dose-dependent increase in the apoptosis rate of CNE2 cells compared to that of the control cells. And the apoptosis rate in all the treatments was all significantly higher than that in the control (*P* < 0.05), as well as higher than that in the corresponding HUV-C. The results demonstrated that HI-3 induced apoptosis of CNE2 cells after 48 h exposure (Figure 5-1); 5-3)). Same to the fluorescence microscope results, HI-3 exposure did not resulted in significantly improved apoptosis rate in the HUV-C (Figure 5-2); 5-3)).

**Figure 5.**
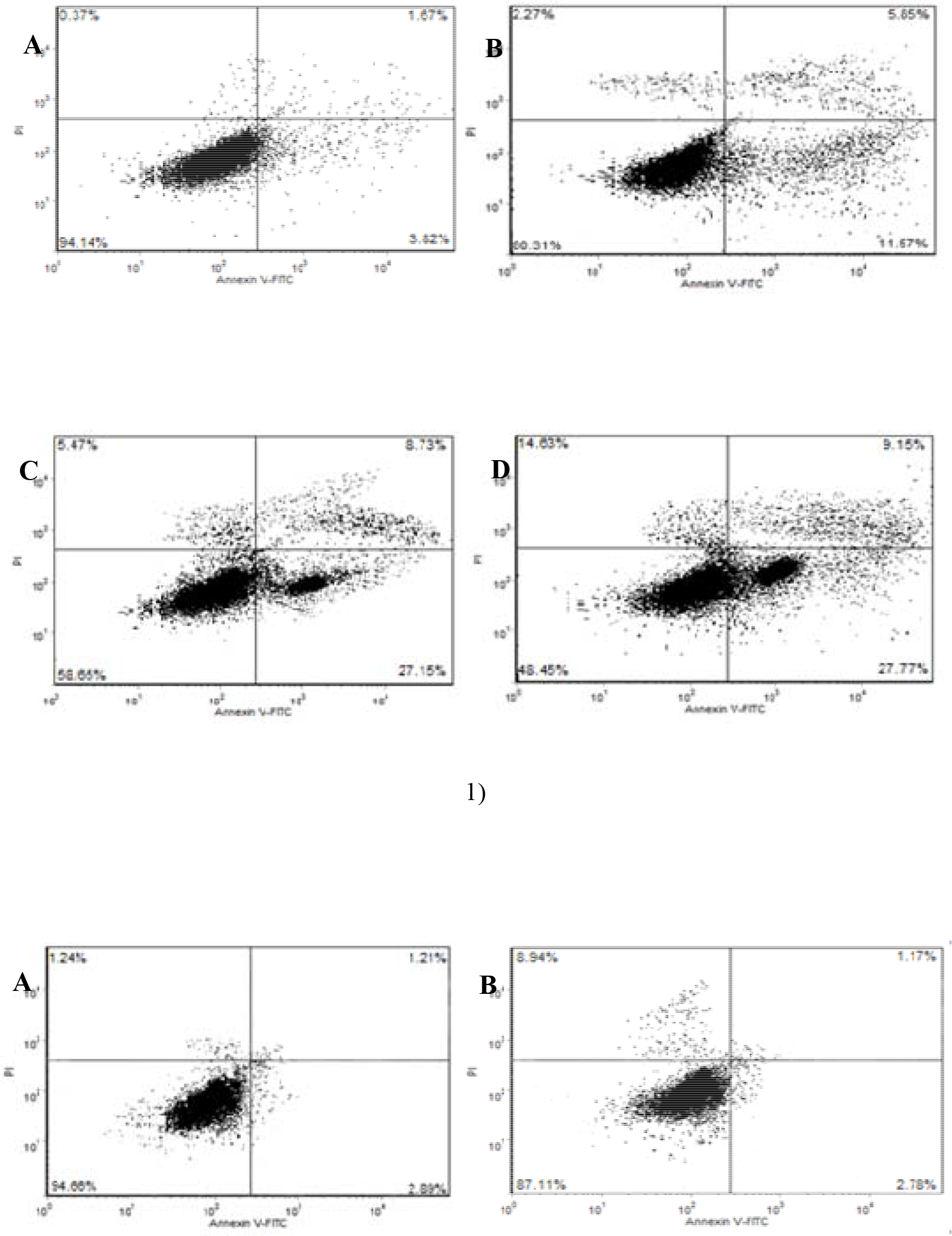
Apoptosis rates of CNE2 cells and HUV-C after treated with HI-3. Note:

1. CNE2 cells A: Negative control; B: 40 μg/ml; C: 80μg/ml; D: 160 μg/ml
2. HUV-C A: Negative control; B: 40 μg/ml; C: 80μg/ml; D: 160 μg/ml
3. Apoptosis rate of HUV-C and CNE2 cells after treated with HI-3 for 48 h. ^#^ *P* < 0.05 means the apoptosis of CNE2 cells were significantly higher when compared with the CNE2 cells in the negative group. △*P* < 0.05 means there was significant difference between CNE2 cells and HUV-C group.

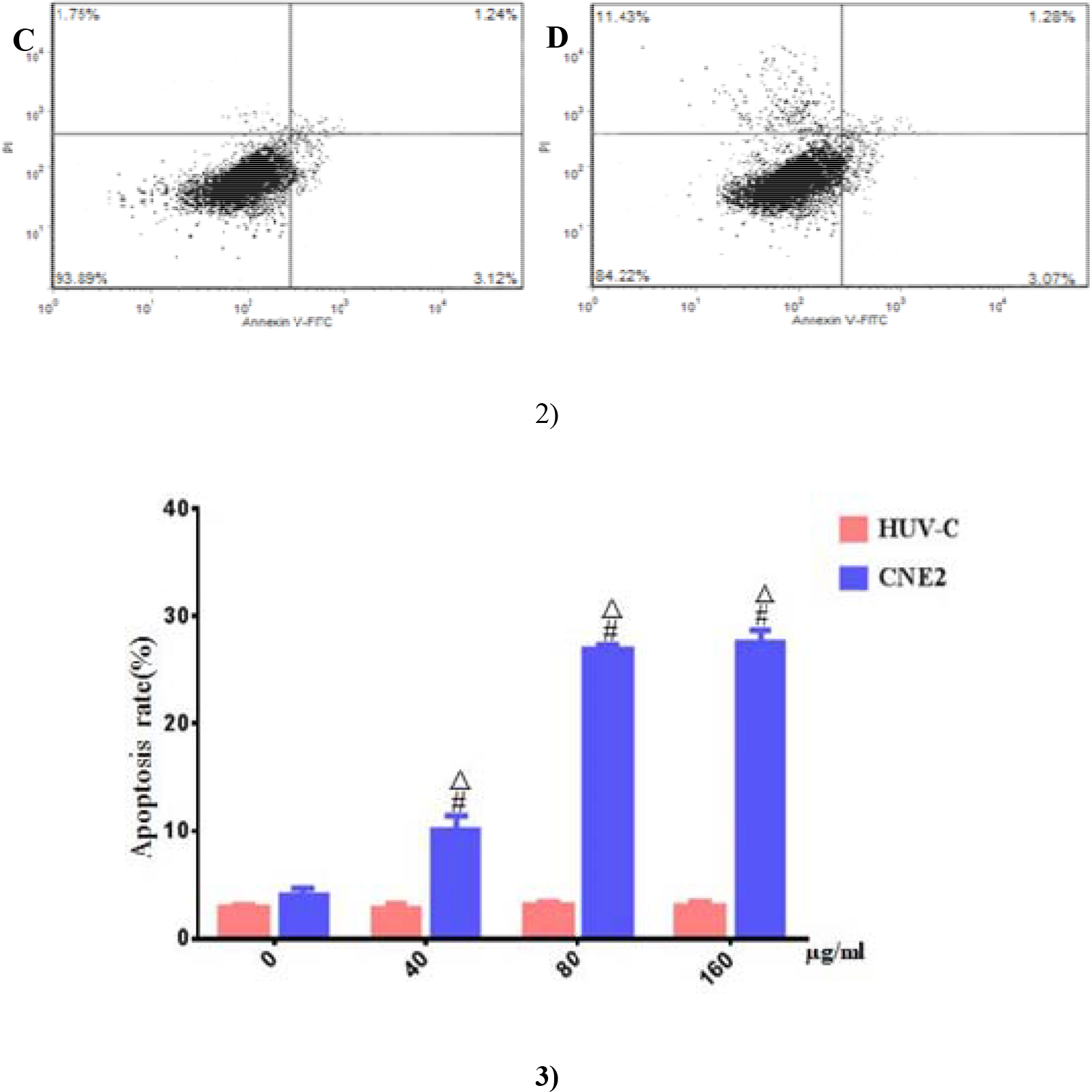

### 3.6 HI-3 inhibited the Migration of CNE2 cells

After making a wound with a pipette tip, it was observed that HI-3 effectively inhibited the migration of CNE2 cells in time-dependent manners when compared to the untreated control cells. The migration rate of CNE2 cells exposed to 160 μg/ml HI-3, which was 24.43 ± 0.47%, 61.5 ± 0.04% and 80 ± 0.33%, respectively, were all significantly lower than that of cells in the control (50.3 2± 0.24%, 91.17 ± 0.15% and 100%) (*P* < 0.05) (Figure 6).

**Figure 6.**
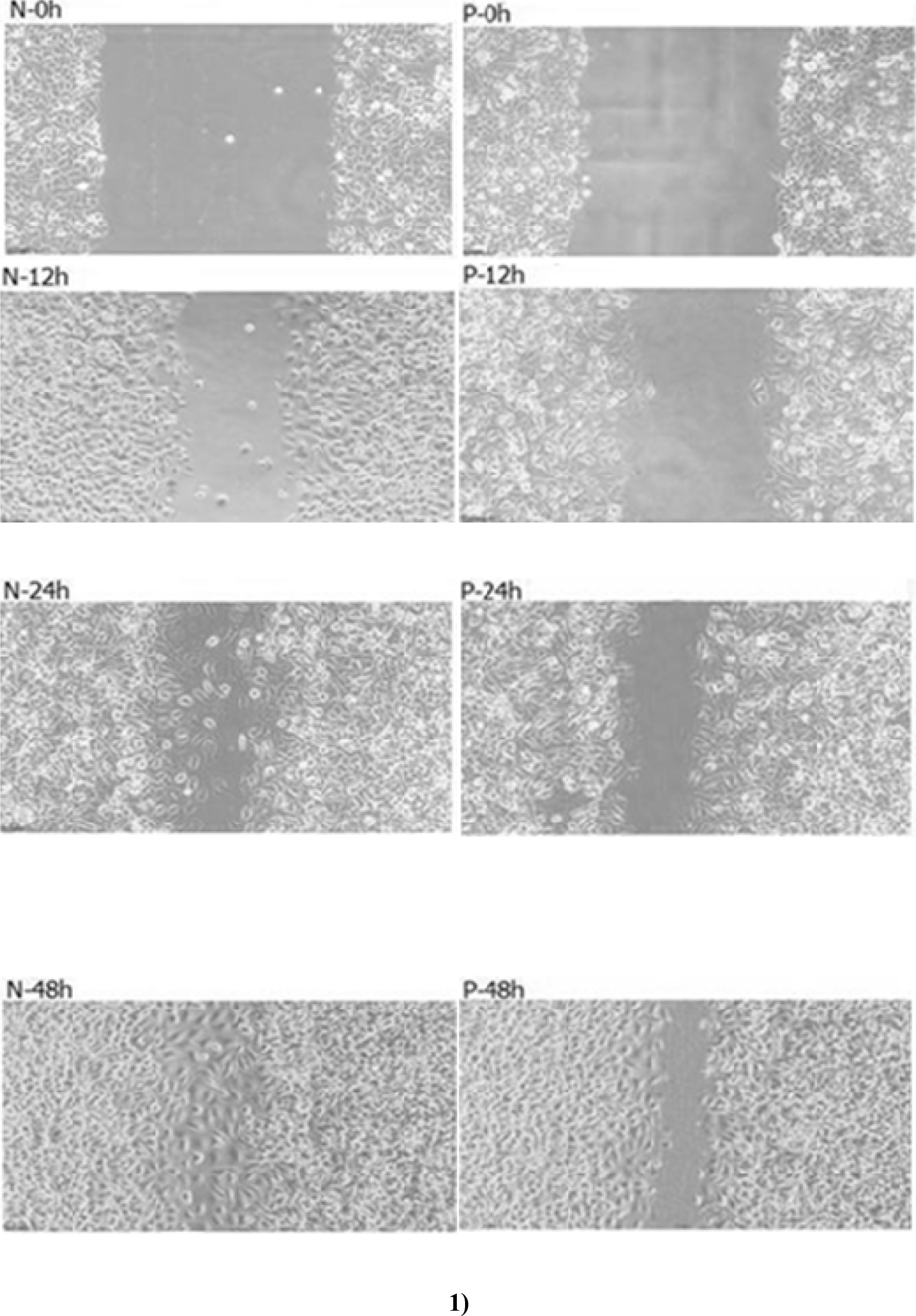
Effects of 160 μg/ml of HI-3 on the migration of CNE2 cells (100×) 1. Morphology observation, N: Negative control group; P: 160 μg/ml of HI-3 group
2. Migration rates of CNE2 cells

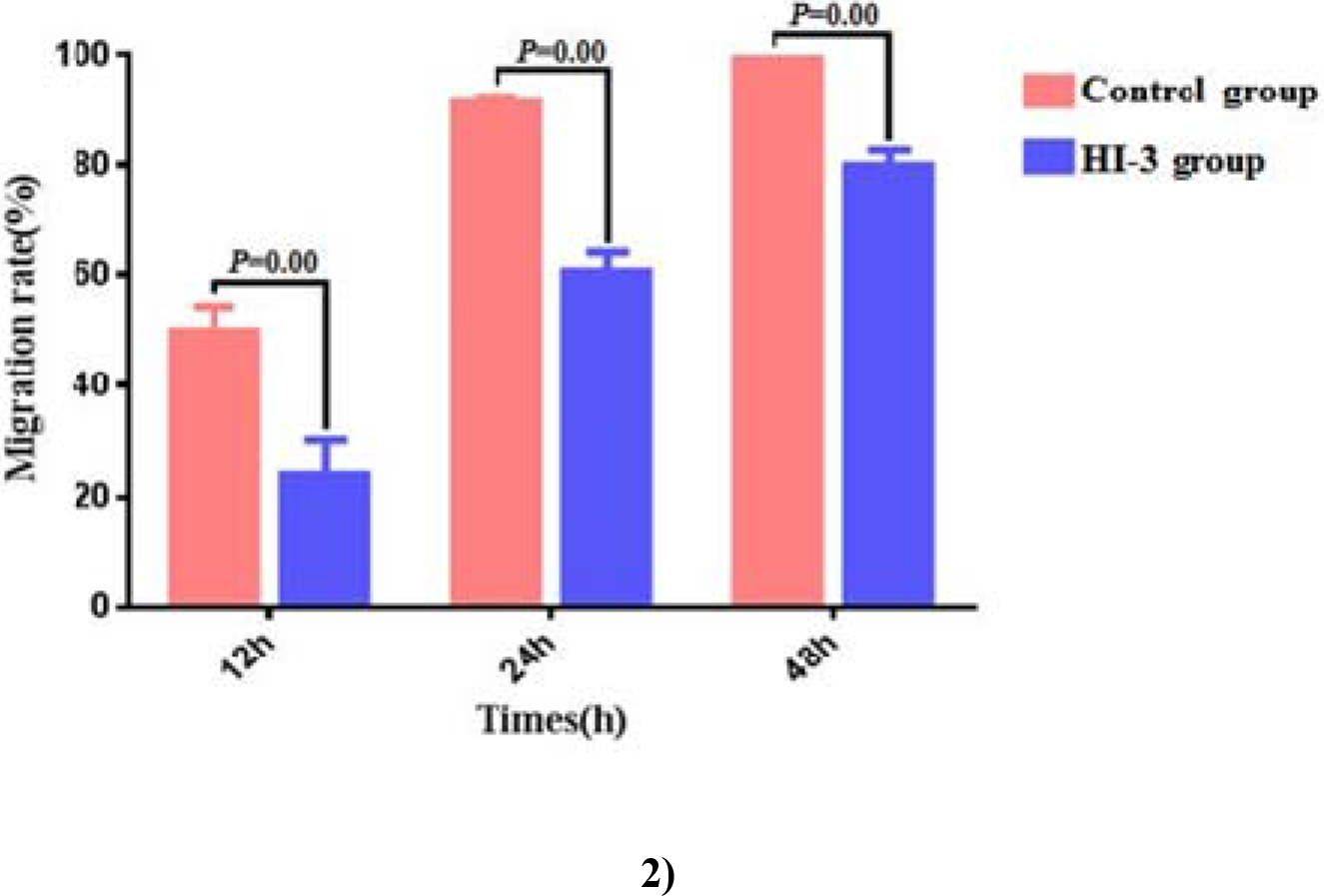

### 3.7 Effect of HI-3 on telomerase hTERT gene expression in CNE2 cells

As showed in Figure 14, the brightness of 28S band is about 2 times of that of 18S band, and no tailing phenomenon occurred, indicating that the RNA quality is qualified without degradation and DNA contamination. After HI-3 treatment for 48 h, the expression of hTERT mRNA was significantly down-regulated in the CNE2 cells compared with that in the control (*P* < 0.05) (Figure 7-1); Table 3). Simultaneously, telomerase hTERT protein expression also significantly decreased. The expression of hTERT protein in the control cells (1.19 ± 0.21) was significantly higher than that in the HI-3 treated cells (0.63 ± 0.12) by gray value analysis (*P* < 0.05) (Figure 7-2), 7-3)).

**Figure 7.**
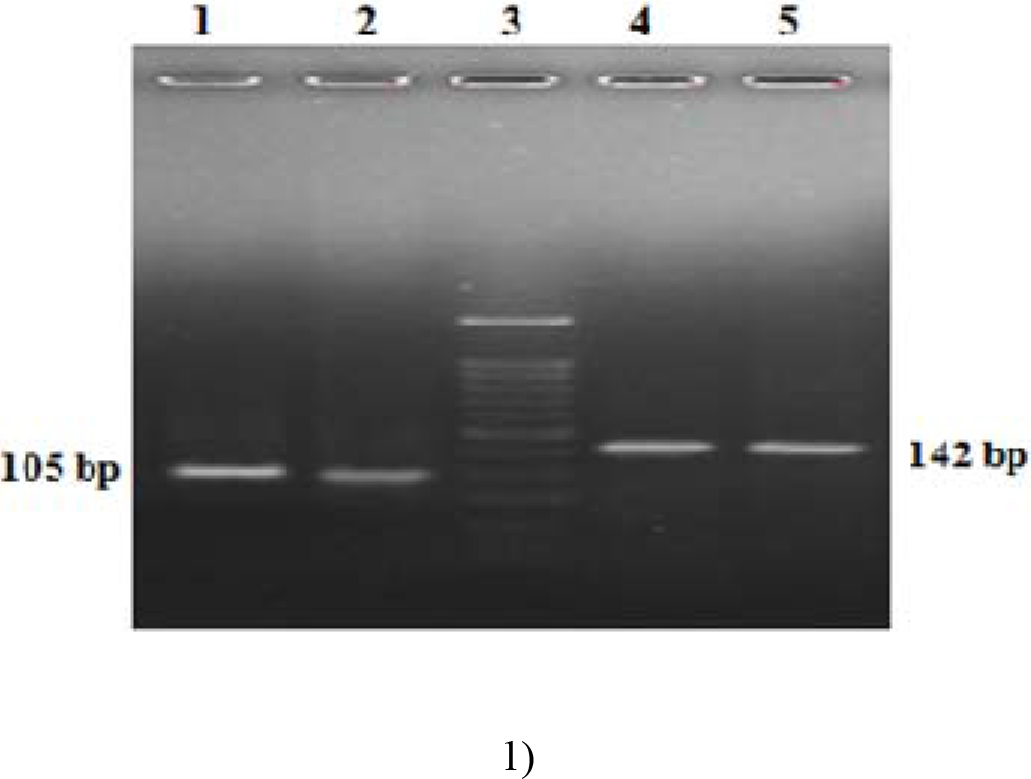
Expression of telomerase hTERT and GAPDH in CNE2 cells after treated with HI-3. 1. RT-PCR analysis on hTERT gene expression, Note: 1: Negative control group; 2: 160 μg/ml HI-3 group; 3: Standard nucleic acid molecule; 4, 5: Internal gene GAPDH;
2. Western blotting on hTERT protein expression;
3. Statistics of expression of hTERT protein.

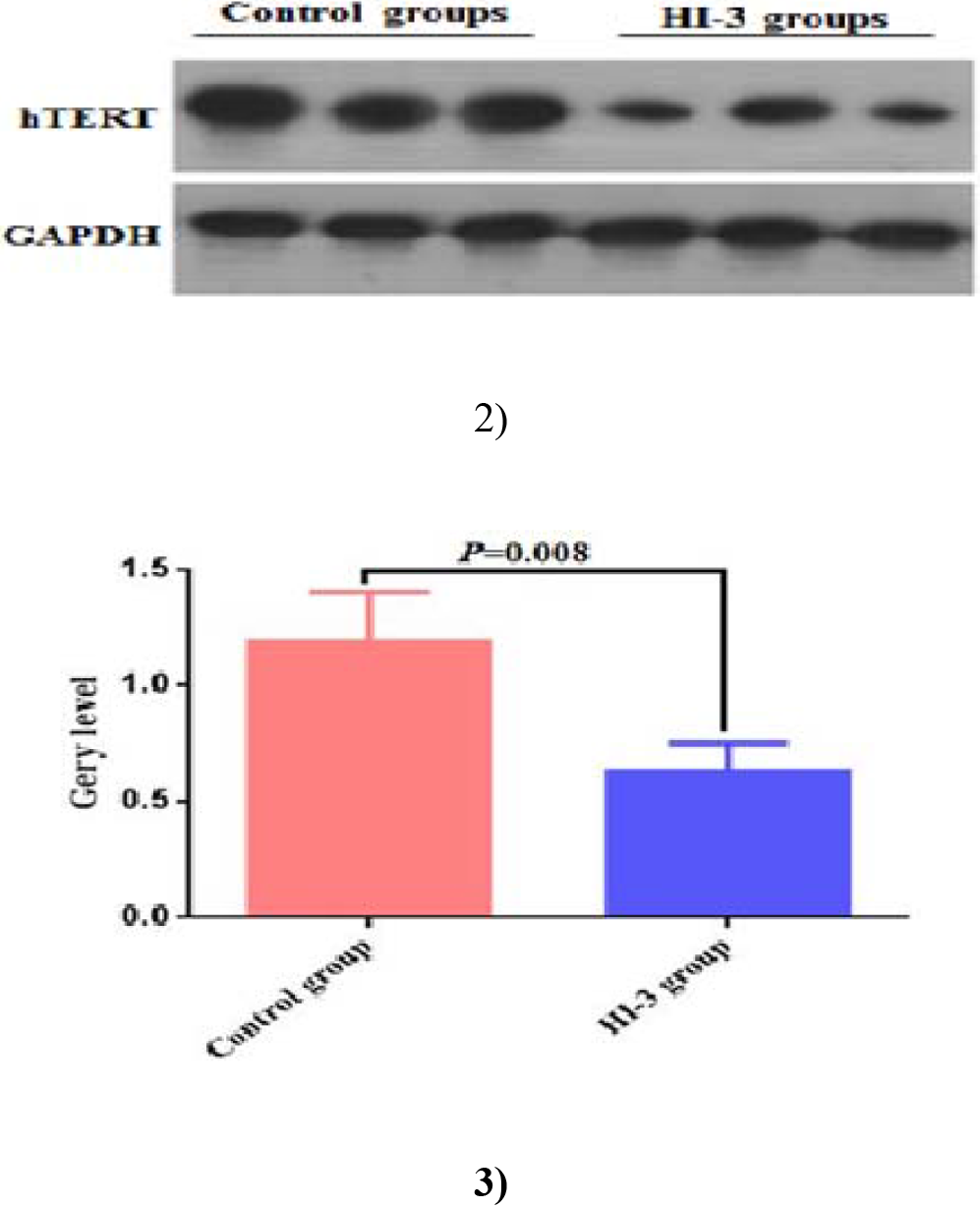

**Table 3.**
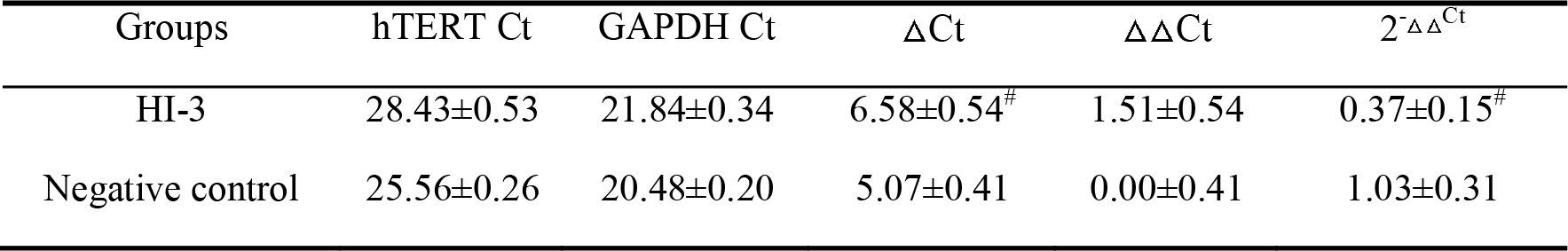
Effect of HI-3 on the expression of telomerase hTERT gene ( n=6,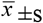) Note: ^#^ *P* < 0.05 indicated that there were significant differences in expression levels between HI-3 group and negative control group.

| Groups | hTERT Ct | GAPDH Ct | ΔCt | ΔΔCt | 2<sup>-ΔΔCt</sup> |
| --- | --- | --- | --- | --- | --- |
| HI-3 | 28.43±0.53 | 21.84±0.34 | 6.58±0.54<sup>#</sup> | 1.51±0.54 | 0.37±0.15<sup>#</sup> |
| Negative control | 25.56±0.26 | 20.48±0.20 | 5.07±0.41 | 0.00±0.41 | 1.03±0.31 |

## 4 Discussion

In this study, we demonstrated that a small molecule peptide HI-3, purified from the hemolymph of *H. illucens* larvae after acupuncture induction with *Staphylococcus aureus*, showed inhibitory effects on both gram-positive and gram-negative bacteria, as well as on the proliferation of CNE2 cells to some extent. It was also found that HI-3 could induce the apoptosis and inhibit the migration of CNE2 cells. Further studies confirmed that HI-3 could significantly inhibit the expression of hTERT. An interesting observation in this study is that HI-3 had more deleterious effects towards CNE2 cells than toward normal cells. Its antitumor activity towards CNE2 cells was unlike that of chemotherapeutic agents, for they cannot discriminate between normal and cancer cells. These results support our assumption that HI-3 may act on a potential antitumor drug without toxic development.

After acupuncture induction with *Staphylococcus aureus*, three components were separated from the hemolymph of *H. illucens* larvae. However, only HI-3 showed different degrees of antibacterial activities against all selected bacteria, which was consistent with the broad-spectrum antibacterial effects of AMPs reported in the previous studies. Generally, AMPs exert their antibacterial activity through the destruction of the plasma membrane (Lee et al., 2015a; Nguyen et al., 2011). Due to their amphipathic nature, most of AMPs are positively charged and can interact with the negatively charged phospholipid bilayer on the bacterial surface by electrostatic action or with other anions on the membrane to form ion channels on the plasma membrane (Huang et al., 2010). It was found that the AMP attacins
from the silkworm could inhibit the growth of gram-negative bacteria by inhibiting the synthesis of membrane proteins; and moricin, another antibacterial peptide, could exert its antimicrobial activity by increasing membrane permeability (Ravi et al., 2011). Though the plasma membranes of CNE2 cells were disrupted by HI-3 treatment in this study, whether the mechanism of HI-3 against bacteria was through this action still need further study.

AMPs isolated from insects, including melittin, cecropin related peptides and magainins have been shown to exhibit antitumoral activity for cells derived from mammalian tumors (Lee et al., 2015b; Su et al., 2015). In the present study, HI-3 at the concentration of 40-160 μg/ml also exhibited the proliferation inhibition of CNE2 cells in a dose- and time-dependent manner. However, the negative control cells HUV-C were not affected by HI-3 treatment. These results were similar to the study of Wang et al. (2012), which found that the AMP temporin-1CEa have different degrees of inhibition against the growth of various types of tumor cells, while normal human erythrocytes and HUVSMCs cells were not affected. Different studies have attempted to explain the mechanisms that the tumor cells are more sensitive to AMPs treatments (Wang et al., 2016). For the surface of tumor cells membrane holds more negatively charged phosphatidylserine, abnormal expressed polysaccharide proteins and increased sialylation when compared with the normal cells, some investigations have determined that one of its antitumoral mechanisms was through destructing the cell membrane, which was similar to that of the antibacterial activity. For example, Slaninová et al. (2012) demonstrated that AMP from wild-type bee venom exerted the cytotoxic effects on tumor cells through enhancing the permeability of tumor cell membranes. Chang et al. (2011) reported that the AMP tilapia hepcidin(TH)1-5 could bind to the membranes of HepG2 and HeLa tumor cells, breaking the cell membrane through a process similar to that of cytolysis, and eventually lead to cell necrosis. In addition, the toxicity of AMP D-peptide from beetles against tumor cells mainly depended on the negative charge carried by phosphatidylserine in the tumor cell membrane (Iwasaki et al., 2009). In present study, fluorescence intensity in CNE2 cells increased with the increasing concentration of HI-3 in a dose-dependent manner, indicating that the target site of HI-3 might also be the membrane. Furthermore, the increasing apoptosis rates in CNE2 cells after HI-3 treatment might also be related with the gradually improved membrane permeability, though more researches should be investigated.

HI-3 was able to inhibit the migration of CNE2 cells to a certain extent according to our cell scratch assay. Generally, tumor cells have a strong ability of invasion and metastasis. Distal metastasis and subsequent increased malignant proliferative capacity are among the major problems during the oncotherapy. Relevant study revealed that an amphibian AMP Temporin-1CEa could inhibit the invasion and metastasis of melanoma A375 cells whereby regulating the release of metalloproteinase-2 and vascular growth factor (Zhang, 2015). However, few studies have focused on the specific mechanisms concerning the migration of cancer cells treated with AMP. The deep research remains to be explored.

Higher activity telomerase could be detected in malignant proliferative cancer cells. Therefore, telomerase is considered as an important and effective target for anti-tumor therapy (Chen et al., 2016). For example, the growth inhibition of human colon cancer HCT166 cells by AZT (3’-Azido-3’-deoxythymidine) was mediated by inhibiting telomerase activity and hTERT gene expression (Hu and Xu, 2017). Our study also found that hTERT gene expression and hTERT protein expression were significantly decreased after HI-3 treatment, indicating that the down-regulating of telomerase activity was involved in the inhibition of NPC. On the other hand, it was reported that in addition to its function of maintaining intracellular DNA and chromosomes stability, telomerase is also involved in the regulation of mitochondrial function and gene expression during tumorigenesis (Chiodi and Mondell, 2012). For hTERT is the key protein of telomerase synthesis, it might also be involved in the action of telomerase in the occurrence and development of cancer. However, the regulation effect of telomerase and hTERT in the HI-3 induced inhibition of proliferation of CNE2 cells still need further investigation. And whether the mitochondrial pathway was induced in the HI-3 action also remains to be explored.

From these results, we concluded that HI-3 has the potential for development as a new type of antitumoral agent, for HI-3 is less toxic to normal cells. It was confirmed that HI-3 might exert its antitumoral effect through down-regulating the expression of hTERT. However, the specific mechanism involved in the HI-3 still need deep investigations.

## Acknowledgements

We thank Prof. Guren Zhang for comments that greatly improved the manuscript, and we thank the “anonymous” reviewers for their constructive suggestions.

## Competing interests

The authors have declared that no competing interests exist.

## Author contributions

Zhong Tian and Qun Feng performed the experiments, Hongxia Sun and Ye Liao drafted and revised the manuscript, Lianfeng Du and Rui Yang performed the data analyses, Xiaofei Li and Yufeng Yang helped perform the analysis with constructive discussions, and Qiang Xia designed the experiments.

## Funding

This work was supported by the grants from The Major Research Project of the Innovation Group of the Education Department of Guizhou Province (KY characters [2016]037 in Guizhou Province); The National Nature Sciences Fund (31260528); and Guizhou Science and Technology Fund (KY characters [2009]2295 in Guizhou Province).

